# The contraction in inverted repeat regions in the complete plastome sequence of *Cressa cretica* L

**DOI:** 10.1101/271221

**Authors:** Pritesh P. Bhatt, Vrinda S. Thaker

## Abstract

Plastome studies have been the focus of research in plant molecular evolution and systematics. *C. cretica* L. (Convolvulaceae) is a halophyte, habitat in the ecologically challenged area with high salinity and drought. The complete physical map of plastome revealed that it is 141,419bp long, circular molecule. It contains typical quadripartite structure of large single copy region (LSC 94,808bp), small single copy region (SSC 32,527bp) separated by a pair of inverted repeat regions (IRs 7042bp). This plastome is compared with the complete plastomes of other members of Convolvulaceae showed notable distinctions. An exceptional shift in IRs to SC regions is experienced in *C. cretica* led to many genes shift in both SC regions and contraction in IRs. The size of IRs reduced to 2 to 4 times as compared to those of the Convolvulaceae members studied. The shifted IRs regions showed remarkable variation in nucleotides patterns. Further, the shift was from the IR boundaries and in between the IR regions led to segment IRs. It is concluded that the shift in IRs may be the strategic move for adaptation in the harsh environment.

## Introduction

The higher plant cells, three compartments contain their own DNA of which the chloroplast harbors the least complex genetic material. The chloroplast genome (plastome) exists as a covalently closed, double stranded circle ranging in size from 85-217 kb, but most common land plants contain 115-164kb. With few exceptions, higher plant chloroplast DNA contains two inverted, exact repeats of 20 to 30 kb which are separated by a small and a large single copy region. Thus a typical circular plastome has a quadripartite structure and exhibits highly conserved gene order and contents (Jansen et al. 2005; Wicke et al. 2011). The number of genes ranged from 63-209 among various lineages, a common pattern of most plastomes showed average 110-130 which includes protein coding, rRNA, and tRNA genes. However, large-scale genome rearrangement and gene loss have been identified in several angiosperm lineages (Wolfe et al. 1992; Lee et al. 2007).

The key important feature of chloroplast DNA is the presence of two large inverted repeats which reduced the rate of new recombination, mutations and promote DNA repair mechanism hence slow down the rate of evolution. Dynamic changes in IRs as expansion/contraction play a decisive role in plastome evolution. In number of studies such as Apioideae (Plunkett and Downie 2000; Downie and Jansen 2015), monocots (Wang et al. 2008), ferns (Wolf et al. 2010) and Pinaceae (Lin et al. 2010) reported that IRs contributes in increase/decrease in size of plastomes and gene rearrangement (Magee et al. 2010).

Investigation of genome size changes in various organisms found correlations with cytological, physiological, and ecological characters (Jockusch 1997; Gregory 2002; Knight and Ackerly 2002), even within single species (Bennett and Leitch 1995; Nevo 2001). *Cressa cretica* (Convolvulaceae) is a perennial halophyte which dominates both inland and coastal marshes. *C. cretica* seeds can germinate in up to 850mM NaCl and it can tolerate up to 950mM NaCl which is one of the highest concentrations (Priyashree et al. 2010). However, little is known about a molecular aspect of this plant. In this study, it is collected from desert of Little Runn of Kutch (Western India). Kutch desert is a unique ecosystem, with an admixture of saline, marshy and coastal desert where water and soil are extremely saline. It also demonstrates high temperature (~48° C) in hot summer and as low as 10° C in winter cold waves. Rainfall is also not adequate (~320 mm). In this harsh environment, this area harbors it own exceptional flora, with some endemic and the species of high conservation significance at national and international levels.

In this paper, we investigate the organization and evolution of cp genome of *Cressa cretica* and it is compared with the available complete plastomes of other members in the Convolvulaceae in the GenBank. The unusual features of IR contraction in the genomes of *C. cretica* are described which provide valuable insights into cpDNA evolution. The comparisons with other members of Convolvulaceae identify multiple inversions, gene duplications, IR contraction, gene and intron losses. Gene relocation caused by IR loss in SC region is a notable feature of the cp genomes of the *Cressa cretica*.

## Results and Discussion

### Genome content and organization

The size of the plastome of *C. cretica* is 141,419bp with typical quadripartite structure, including a LSC region of 94,808bp and SSC region of 32527bp separated by a pair of identical IRs of 7042bp each (Fig.1). The total plastome size is consistent with those from other angiosperms which range from 85 to 176 kb. The complete plastome of *C. cretica* is slightly shorter than that of *Ipomoea* species (161-162 kb) but larger than those of *Cuscuta* species (85-125 kb) which all belong to the Convolvulacease (Table-1). The length of LSC region in *C. cretica* was slightly higher to *Ipomoea* species but much larger than that of *Cuscuta gronovii* and *C. obtusiflora*. In *C. exaltata* and *C.reflexa*, LSC region recorded lower values than that of *C. cretica* and *Ipomoea* species (Table-1). In contrast to other members of Convolvulacease, *C. cretica* showed extended SSC region from both ends of IRs, the largest size i.e.32.527 kb (Table-1, Fig.1). Further, in *C. cretica*, a total number of the protein coding genes were 82, total 27 tRNA coding genes for 21 different tRNAs, and 8 rRNA genes for 4 rRNA were observed. It has near to equivalent numbers of protein coding, tRNA and rRNA genes to *Ipomoea* species (Table-2).

**Table 1.** Comparative analysis of plastome data in *Cressa cretica* and other members of Convolvulaceae

| Species | Size<br>(kb) | IRa region | IRb region | LSC | SSC |
| --- | --- | --- | --- | --- | --- |
| *Cressa cretica* (Cc) | 141.419 | 7.042 | 7.042 | 94.808 | 32.527 |
| *Cuscuta exaltata* (Ce) | 125.373 | 16.701 | 16.701 | 82.721 | 9.250 |
| *Cuscuta gronovii* (Cg) | 86.744 | 14.354 | 14.354 | 50.973 | 7.063 |
| *Cuscuta obtusiflora* (co) | 85.286 | 14.131 | 14.131 | 50.207 | 6.817 |
| *Cuscuta reflexa* (Cr) | 121.521 | 16.741 | 16.741 | 79.468 | 8.571 |
| *Ipomoea batatas* (Ib) | 161.303 | 20.692 | 20.692 | 87.823 | 32.089 |
| *Ipomoea nil* (In) | 161.897 | 30.847 | 30.847 | 88.117 | 12.086 |
| *Ipomoea purpurea* (Ip) | 162.046 | 30.882 | 30.882 | 88.172 | 12.110 |

**Fig. 1.**
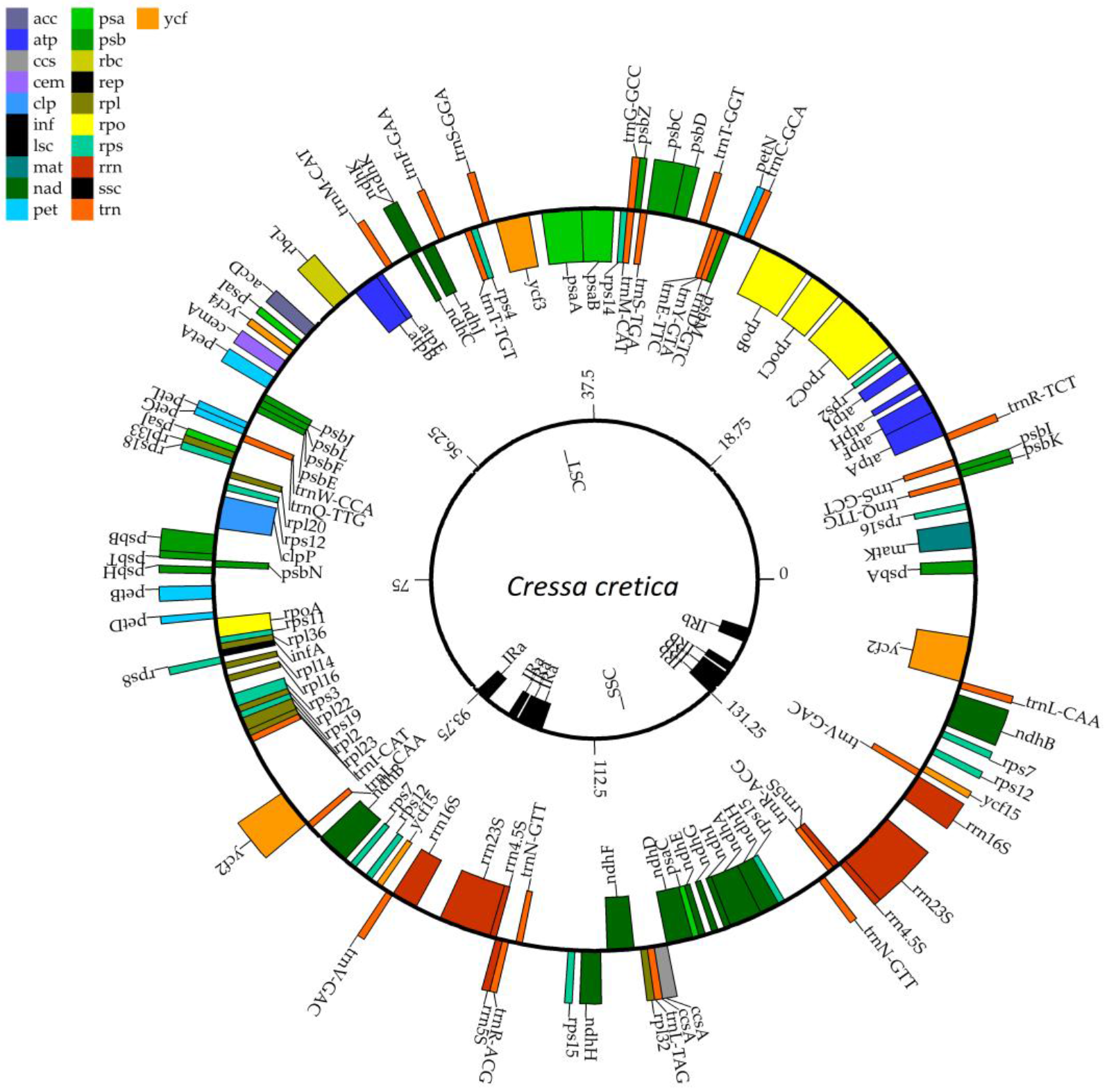
Genome map of *Cressa cretica* chloroplast prepared using CPGAVAS server (Liu et al. 2012). Genes shown in the clockwise direction on outside of the structure is transcribed in anti clockwise direction while genes shown on the inside of the circle are transcribed in the clockwise direction. Legend indicates the functional group to which each gene belongs

### GC content

Plastome GC content is highly conserved in land plants and is typically in the range of 30-40%, with GC content being lower in non-coding regions than in coding regions (Bock 2007). In Convolvulaceae plastomes, an overall higher range of GC content was observed 37.5-38.6% (Table-2). In all the members of Convolvulaceae, higher GC content observed in rRNA and tRNA genes (50-55%) than in protein coding genes (37.5-39%). The overall G+C content was slightly higher in *C.cretica* than other members of Convolvulaceae.

**Table 2.** Comparative studies on genomic data of *C. Cretica* and other members of Convolvulaceae

| Feature | *Cressa cretica* | *Ipomoea nil* | *Ipomoea purpurea* | *Ipomoea batatas* | *Cuscuta reflexa* | *Cuscuta obtusiflora* | *Cuscuta gronovii* | *Cuscuta exaltata* |
| --- | --- | --- | --- | --- | --- | --- | --- | --- |
| Entire chloroplast genome size (bp) | 141419 | 161897 | 162046 | 161303 | 121521 | 85286 | 86744 | 125373 |
| No. of genes | 121 | 132 | 131 | 141 | 113 | 98 | 98 | 117 |
| No. of Proteins | 82 | 87 | 85 | 93 | 69 | 61 | 62 | 67 |
| No. of tRNA | 27 | 38 | 38 | 40 | 35 | 29 | 28 | 35 |
| No. of rRNA | 8 | 8 | 8 | 8 | 8 | 8 | 8 | 8 |
| No. of genes with introns | 10 | 12 | 10 | 10 | 9 | 2 | 1 | 6 |
| GC content (%) | 38.6 | 37.47 | 37.48 | 37.59 | 38.22 | 37.84 | 37.72 | 38.12 |
| GC content for gene sequences (%) | 38.9 | 38 | 38.1 | 38.1 | 38.7 | 37.4 | 37.4 | 38.4 |
| GC content for coding sequences (%) | 39.1 | 38.4 | 38.5 | 38.5 | 39.1 | 37.5 | 37.5 | 38.8 |
| GC content for rRNA genes (%) | 54.5 | 54.9 | 54.9 | 54.7 | 55.2 | 54.7 | 54.7 | 55.3 |
| GC content for tRNA genes (%) | 52.7 | 49.6 | 51.7 | 51.7 | 52.8 | 50.2 | 51.2 | 52.7 |
| NCBI accession numbers | NC_035516 | NC_031159 | NC_009808 | NC_026703 | NC_009766 | NC_009949 | NC_009765 | NC_009963 |

### IR loss

The outstanding structural feature of plastome of land plants studied so far is a pair of large inverted identical repeat (IRs) sequence, which varying in length from 15-30kb in different species (Raubeson and Jansen 2005; Bock 2007). In Convolvulacease family, considerable length variation exists between the plastomes; with the smallest IR (7042bp) is observed in *C. cretica* less than one-forth size of that in *Ipomoea purpurea* and *I.nil*, less than two and half size of *I. batatas* while nearly less than half size of IR in *Cuscuta* species (Table-1). Slight variability in the size of land plan IRs is due to expansion and contraction of IR boundaries into single copy regions (Raubeson and Jansen 2005; Bock 2007); however, the extent of IR contraction in *C. cretica* is unprecedented. Here, the region is more than 2.5 to 5 times smaller than the typical land plants IR including the members of Convolvulacease compared (Table-1).

Notably, IR regions divided into four segments; IRaI to IRaIV and IRbI to IRbIV (Fig.2a). The loss of base pairs in IRa and IRb also observed significantly different. Between IRaI to IRaIV, the loss observed for 17, 271 and 3449 bp, respectively; while in IRbI-IRbIV, it was 37,270 and 3737bp, respectively. Thus a total loss of 3737 (in IRa) and 3796 (in IRb) was observed, with a difference of 59bp loss at IRa site (Fig.2a). In LSC, IR boundary shift was evident with *trnL-CAA* and *ycf2* (Fig.2b). Further, the genes shifted in SSC from IRa and IRb also aligned showed a significant difference in intergenic as well as in coding regions of *rps7, rps12, ycf15* and *ndhH* genes (Fig.2c). Pairwise alignment of nucleotide sequences of IRa and IRb regions, shifted in either LSC or SSC regions, showed significant nucleotide variations (Table-3). More polymorphic sites were observed in LSC (225) than in that of SSC (16) and indel sites were also higher in LSC (812) than in SSC (44). These data suggest that the loss of IR region is more towards the LSC region. LSC/IR and SSC/IR boundaries are sometimes regarded as an index of chloroplast evolution. As observed in other plastome studies, the IR contraction has led to changes in the structure of the chloroplast genome, contributing to the formation of pseudogenes (Saski et al. 2005; Zhang et al. 2013; Luo et al. 2014). In this study, we have observed *rps7* and *ycf2* on IRb shift and *ndhH* on IRa shift as pseudogenes.

**Fig. 2.**
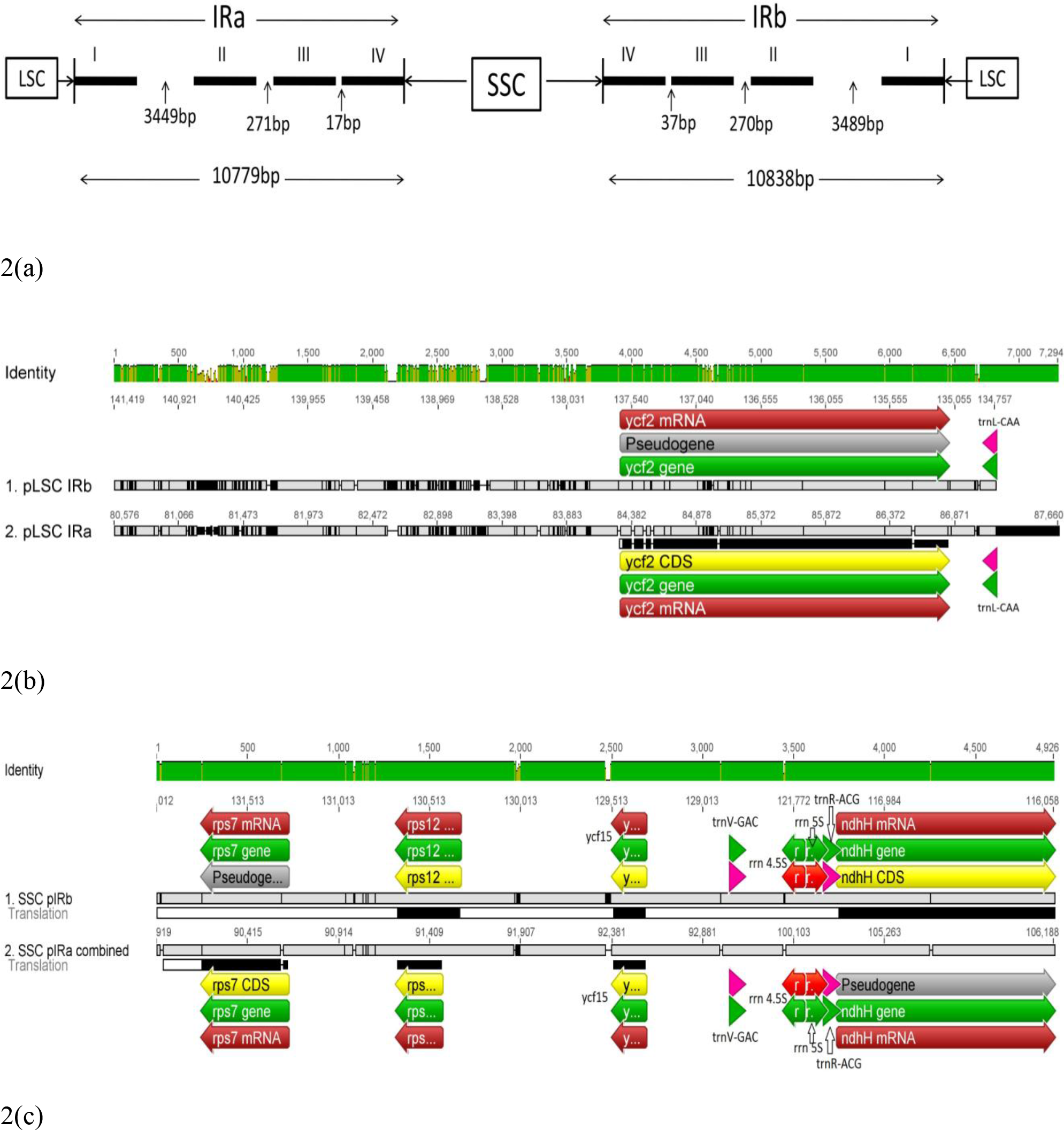
(a). LSC/IR and SSC/IR boundaries on *Cressa cretica* plastome. IRaI-IV and IRbI-IV shows segmented inter repeat regions, with bp gap in between. (b). Gene shift in LSC/IR boundaries. Broken lines indicate indels on the shifted sequences whereas solid lines indicate complete gene frame. (c). Gene shift in SSC/IR boundaries; Otherwise as per fig.2b.

**Table 3.**
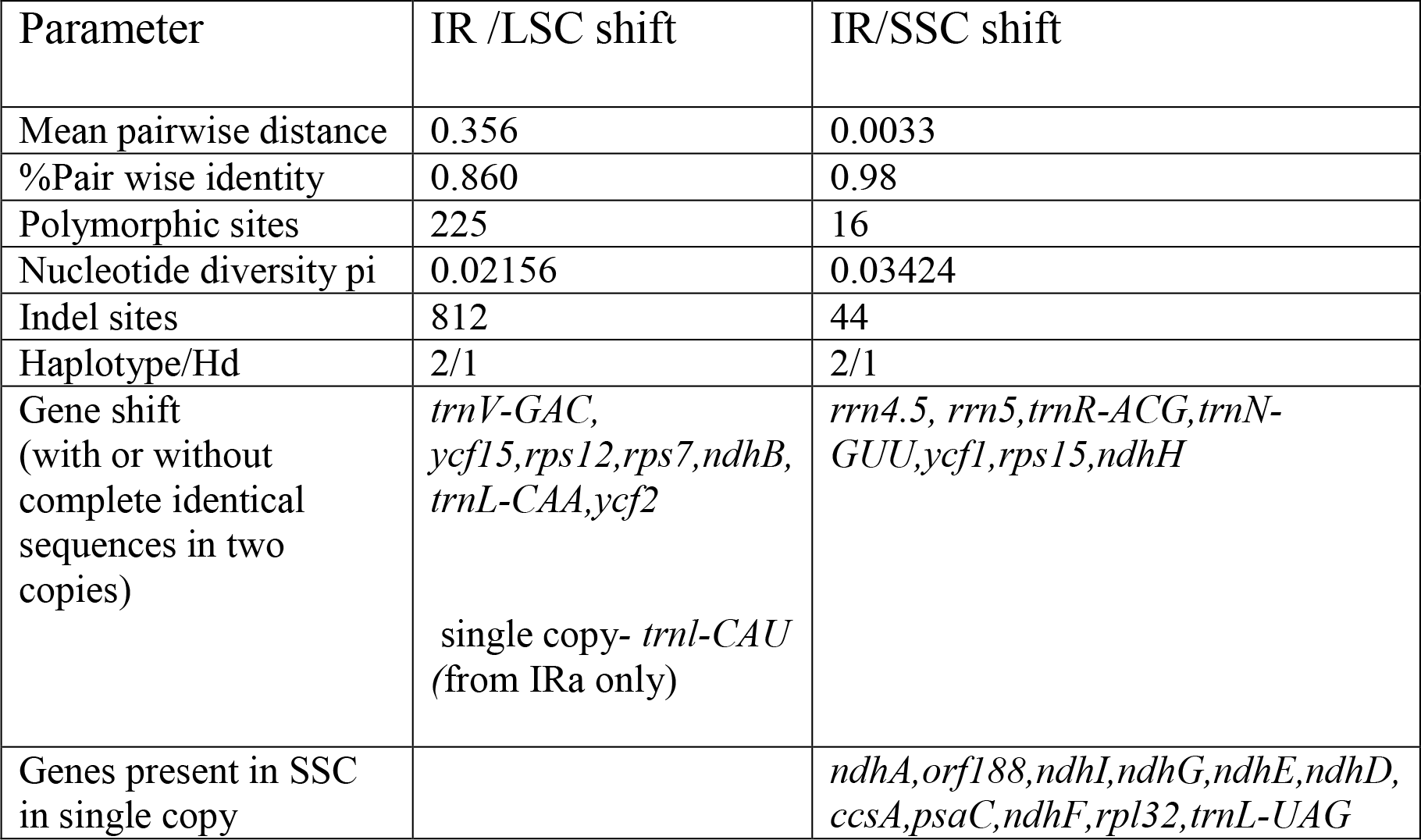
Comparative studies on nucleotide diversity from extended LSC and extended SSC regions due to IR loss in *C. cretica*

### Gene rearrangement

IR contains a core rRNA and tRNA cluster, this includes four rRNA genes for 4.5S,5S,16S and 23Sand five tRNA genes encoding trnA-U-GAC, trnI_GAU, trnN-GUU, trnr-ACG and trnV-GAC (Jansen et al. 2005). In addition, IR region also contains a variety of other genes as a result of lineage-specific expansion and contractions (Zhu et al. 2016). However, in *C. cretica*, each IR includes only 2 copies of *trnl-GAU*, two copies of *trnA-UGC, orf56, orf42, rrn16, rrn23, ndhB* and *ycf68*: remaining IR region shifted in SC region. In SSC, genes for *rps7, rps12, ycf15, trn_VGAC, rrn4.5, rrn5, trnR-ACG, trnN-GUU, ycf1* and *ndhH* were observed in two copies, however, significant substitutions among the copy of these genes were observed. One copy of *rps7* and *ndhH* observed as pseudogenes (Fig.S1). In general SSC region contains the single copy of NADH oxidoreductase genes *ndhA,ndhD, ndhE, ndhF, ndhG, and ndhI* along with genes for photosystem I-*psaC*, ribosomal protein small subunit *rps7, rps15* and ribosomal protein large subunit and *rpl32* On the other hand, in LSC shifted region of IRs *trnL-CAA* and*ycf2* were observed in two copies with remarkable nucleotide variations (Table-3, Fig.2b). The shifted regions showed nucleotide variations in either gene sequences (*rps7, ndhH and ycf2*) or the shift was due to changes in intergenic regions (Fig.2&4 Table-4). *ycf2* found reduced in size like pseudo alternative of *ycf2* found in *Cuscuta reflexa*. Other than highly diverse *ycf2* one copy is pseudo in *Cressa*. In *Cressa ycf2* shifted in SSC is 2502bp in size while of LSC (pseudo) is 2498bp in size. *ycf2* of other convolvulaceae members are 6606bp of *Ipomoea nill*, 6627bp of *I.batata*, 6594bp of *I. purpurea*, 6717bp of *Cuscuta exaltata*, 5415bp of *C.gronovii,*, 5394bp of *C.obtusiflora* and 6012bp of *C.reflexa* (Fig. 3). In all three shifted genes from IR of *Cressa* shows strong node support for phylogenetic relationship (Fig.S1) by giving 100% bootstrap value while poor bootstrap observed for *Ipomoea nill*. To further confirm evolutionary pressure for these three shifted genes Ka/Ks ratio was calculated and values obtained are 3.56 for ycf2, 3.62 for rps7 and 3.41 for ndhH. Thus >1 value of Ka/Ks confirms mutation biases among these genes and shows positive selection pressure (Yang 1998; Chen et al. 2017) on these protein coding genes (S Table-S1).

**Fig. 3.**
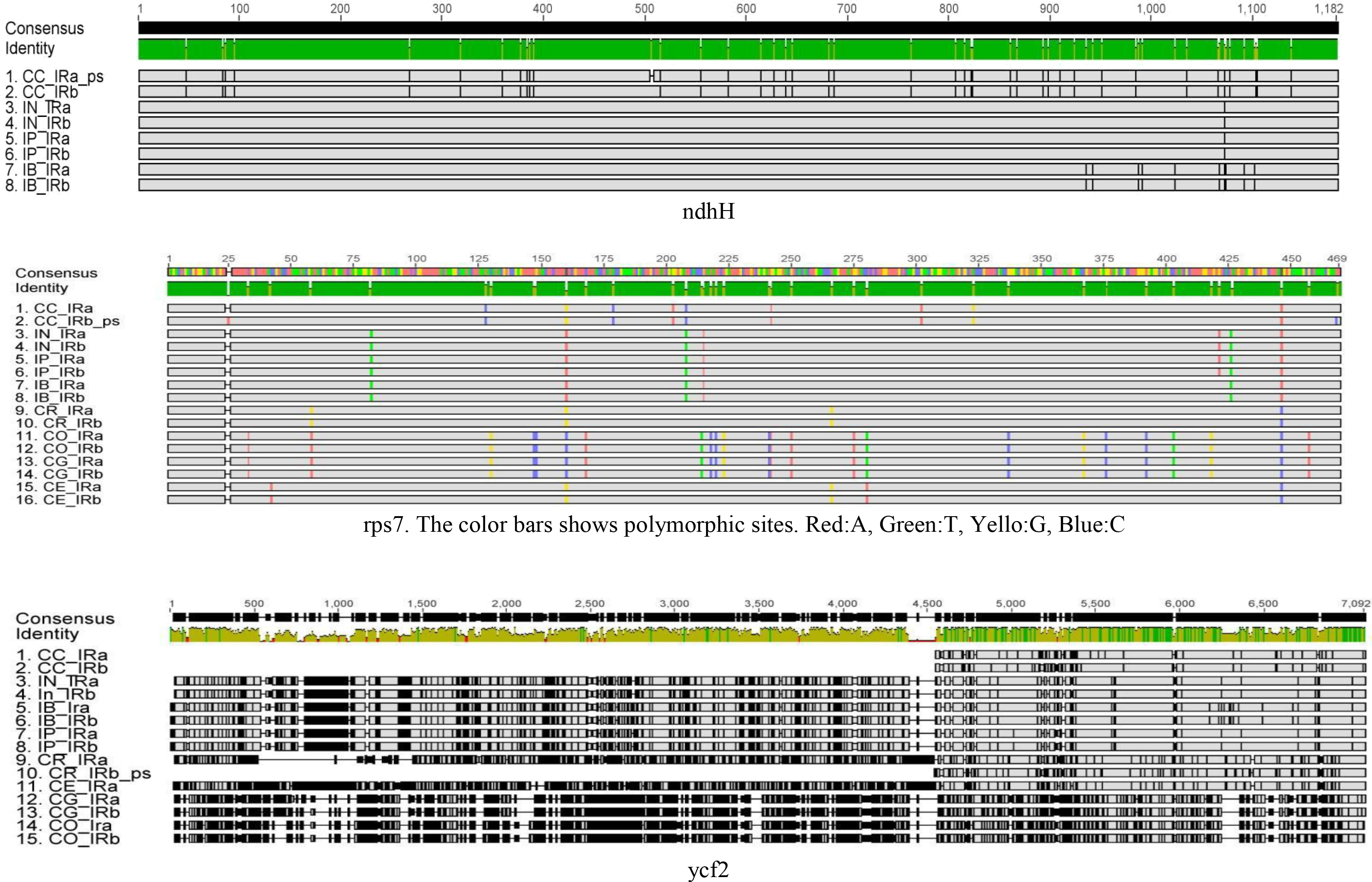
(Comparative alignment of shifted genes of *C. cretica* with other members of Convolvulaceae. Bars in the sequence alignment represent variations in nucleotides between and within the gene sequences. Other details are as per Table-1.

**Table 4.** Variations in number of the genes with introns in Convolvulaceae

| Gene | *Cressa cretica* | *Ipomoea nil* | *Ipomoea purpurea* | *Ipomoea batatas* | *Cuscuta reflexa* | *Cuscuta obtusiflora* | *Cuscuta gronovii* | *Cuscuta exaltata* |
| --- | --- | --- | --- | --- | --- | --- | --- | --- |
| *matK* | - | - | - | - | 1 | - | - | - |
| *atpF* | 1 | 1 | 1 | 1 | 1 |  | - | 1 |
| *rpoC1* | 1 | 1 | 1 | 1 | 1 |  | - | 1 |
| *ycf3* | 2 | 2 | 2 | 2 | 2 |  | - | 2 |
| *ndhJ* | 1 | - | - | - | - | - | - | - |
| *accD* | 1 | 2 | 2 | 2 | 1 | 2 | - | - |
| *clpP* | 2 | 2 | 2 | 2 | 2 |  | 2 | - |
| *ndhB* | 1 | 1 | 1 | 1 | - |  | - | - |
| *ndhA* | 1 | 1 | 1 | 1 | - |  | - | - |
| *ndhB* | 1 | 1 | 1 | 1 | - |  | - | - |
| *trnF*-AAA | - | 2 | - | - | * *trnA* -TGC-1 | *trnK*-TTT-1 | - | * *trnA* -TGC-1 |
| *trnG*-Tcc | - | 2 | - | - | * *trnA* -TGC-1 |  | - | * *trnA* -TGC-1 |
| *Ycf1* | 1 | 3 | 3 | 3 | * *ycf2* -1 |  | - | - |
| NCBI accession numbers | MF 067398 | NC_031159 | NC_009808 | NC_026703 | NC_009766 | NC_009949 | NC_009765 | NC_009963 |

\*indicate different genes with intron in that plant.

**Fig. 4.**
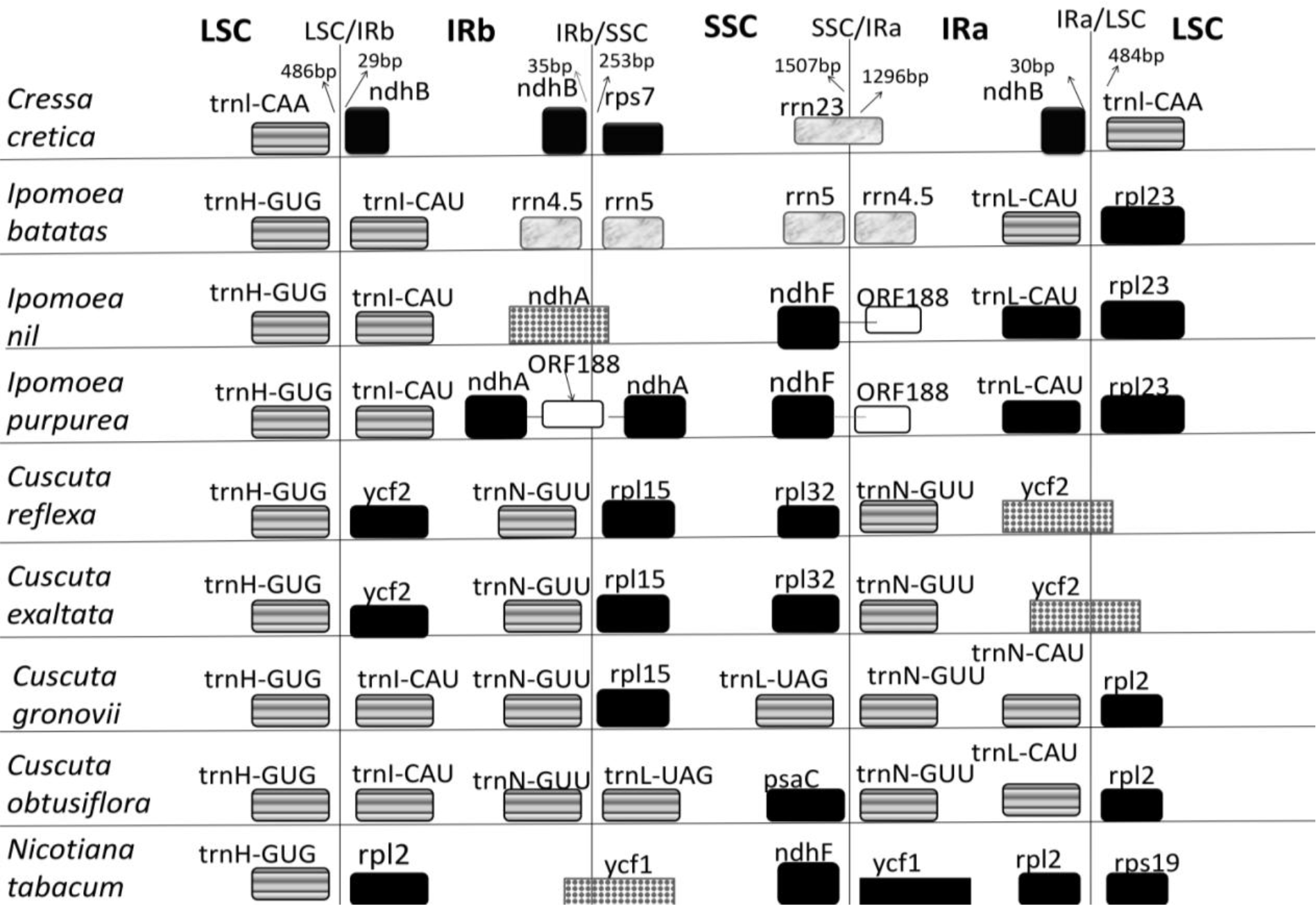
Comparative structures of the LSC /IR and SSC/IR boundaries in Convolvulaceae and *N. tabacum* as out group

The existence of IRs may confer on the plastome by a formation of head-to-head dimmers. IRs represents hotspots for resistance to intra molecular recombination loss and support stabilization of plastome against rearrangement (Wicke et al. 2011). Nevertheless, these potential functional implications do not prevent the variation of IRs. On the other hand, IRs is frequently subjected to expansion, contraction or even complete loss (Palmer et al. 1987; Tsudzuki et al. 1992; Guo et al. 2014). The IR region of other Convolvulaceae plastomes was highly conserved but structure variation still found in IR/SC boundaries (Fig.3). To elucidate the potential contraction of IR regions, we compared the gene variation at the IR/SSC and IR/LSC boundaries regions of eight plastomes. The gene trnH-GUG was observed in all Ipomoea species and *Cuscuta* species including *N. tobacco* as an outgroup, at the junction of LSC/IRb region whereas in *C. cretica* at the LSC/IRb boundary trnI-CAA was observed followed by 468bp extension touching to the IRb boundary. In IRb region, an initial 29bp region was observed before *ndhB*. Two copies of *ndhB* crossed IRb and SSC with an intron sequence in *C.cretica*. Although, the boundary genes also varies between *Ipomoea* species and *Cuscuta* species, compared (Fig.3). The overall location of IR/SC boundaries and genes were found to be varying in Convolvulaceae and *N. tobacco* studied.

In closely related species, IR boundary shift reported being relatively minor resulting in the gain or loss of a small number of genes (Wicke et al. 2014; Downie and Jansen 2015; Wu and Chaw 2015). However, in *C cretica*, exceptional loss of genes from IR to SC regions was observed compared to other members of Convolvulacease. Many plants have lost a major portion of IR or even all, as reported for conifers, many legumes and *Erodium* species (Palmer et al. 1987; Raubeson and Jansen 1992; Guisinger et al. 2011; Guo et al. 2014).

IR boundaries expansion or contraction in SC region up to several hundred bp or by several kb which relocated multiple genes into or out IR has a great impact on plastome DNA evolution (Perry and Wolfe 2002). Although the presence of IRs in plastome is typical, few exception like some algae (Turmel et al. 2005), land plants plastomes lack IR, (Milligan et al. 1989; Cai et al. 2008) indicating that it is not essential feature for plastome function or maintenance (Guisinger et al. 2011). The observation suggests that most genomes that have lost partial or full IR exhibit extensive gene order rearrangements that the presence of the IR plays an important role in stabilizing the plastome (Palmer and Thompson 1982; Hirao et al. 2008). Similarly, lack of IR and rearrangement of plastome in *Trifolium* is reported (Cai et al. 2008).

### Genes with introns

The genes with introns showed significant variations among Convolvulacae. Maximum 12 genes were found in *Ipomoea nil*, followed by 10 in *I.purpurea* and *I.batatas*, 9 in *C.cretica* and *Cuscuta reflexa*, 6 in *C.exaltata* 2 in *C.obtusiflora* and 1 in *C.gronovii* is observed (Table-4). Plastome tRNA gene sequences are generally highly conserved and loss is unusual among angiosperms with some exceptions (Morden and Wolfe 1991; Chumley et al. 2006). In *C. cretica*, loss of an intron in *trnF-AAA* and *trnG-TCC* were observed as compared to *Ipomoea nil*. Further, *C.cretica, ndhJ* showed an intron 27bp, was not found in any other members of Convolvulacease. On the other hand, an intron was lost from ycf1in *C.cretica* and *Cuscuta* species but in *Ipomoea* species, three introns were observed. In general, introns and gene loss were more common in *Cuscuta* species where in *C. gronovii* showed an intron in *clpP* gene only. Thus marked distinction in loss of introns in the genes in Convolvulacease is observed.

Overall gene arrangement on plastomes of Convolvulacease remained nearly uniform. All taxa of *Ipomoea* and *Cuscuta* lack an intact *infA* gene (McNeal et al. 2007). We have also observed 234 bp partial *infA* gene sequences in *C.cretica* support the view that this gene loss probably occurred prior to the divergence of Solanaceae from Convolvulaceae (Schmitz-Linneweber et al. 2001). The loss of *infA* from plastome has been reported in angiosperm evolution (Millen et al. 2001). In addition, *ycf15* gene is lost across Convolvulaceae (McNeal et al. 2007) but present in *N. tobacco*. A third gene *rpl23* is a pseudo gene in *Cuscuta* and function in *Ipomoea* is not clear. The presence of *rpl23* in *C. cretica* needs tests of expression for confirmation.

In sum, an exceptional shift in IRs to SC regions is experienced in *C. cretica*. As a result, many genes shift in either SC region showed remarkable variation in nucleotides patterns. Further, the shift was not only from the IR boundaries but in between the IR regions led to segment IRs. Even the total of pieces of IRs was also 2 to 4 times less than that of any other member of Convolvulaceae compared. *C. cretica* habitat in the environmentally challenged area, the shift in IRs may be the strategic move for adaptation in this harsh environment. However, plastome analysis of some additional plants from this area may help to confirm the conclusion.

## Materials and Methods

### Chloroplast DNA isolation and sequencing

Chloroplast DNA was extracted from leaves of *Cressa cretica* according to Shi et al. 2012 protocol. Chloroplastic DNA was confirmed on 1% agarose gel electrophoresis and concentration was checked. About 10μg of total DNA was used for genome sequencing. Whole genome shotgun sequencing of chloroplast genome was performed using a high throughput ion torrent genome machine with ion torrent server (torrent suite v3.2).

### Genome assembly and annotation

Number and quality of raw reads obtained were evaluated, checked for adapter contamination and average quality score with FastQC v0.11.5 (http://www.bioinformatics.babraham.ac.uk/projects/fastqc/). Reads were quality trimmed using CLC Genomics Workbench v9.5.64 (CLC bio, QIAGEN) with quality score 0.05. Reference guided assembly was performed using CLC (mapping parameters: Mismatch cost=2, Insertion cost=3, Deletion cost=3, length fraction=0.5, similarity=0.8) with the published Convolvulaceae genome of *Ipomoea nil* (NC_031159), as the reference genome. Contigs with >50× sequence depth were used for reference guided assembly. The vote majority conflict resolution mode was used in order to ensure inclusion of only chloroplast specific reads thus avoiding contribution of nuclear and mitochondrial reads to the consensus sequence. Trimmed reads were de novo assembled using CLC. Consensus sequence derived from reference assembly was compared and corrected with de novo assembly. Plastome annotation was performed in DOGMA (Wyman et al. 2004) and CpGAVAS (Liu et al. 2012). All Gene sequences were confirmed by comparing them with available Convolvulaceae genomes and manually corrected. Further tRNA genes were confirmed using tRNAscan-SE 2.0 (Lowe and Eddy 1997).

### Data collection

Complete Convolvulaceae plastome sequences of *Ipomoea batatas* (NC_026703), *Ipomoea nil* (NC_031159), *Ipomoea purpurea* (NC_009808), *Cuscuta reflexa* (NC_009766), *Cuscuta exaltata* (NC_009963), *Cuscuta gronovii* (NC_009765), *Cuscuta obtusiflora* (NC_009949) and *Nicotiana tabacum*- standard (NC_001879) were retrieved from NCBI for comparison and analysis.

### Genome analysis

Gene comparison and graphical views were generated using Mauve plugin in Geneious. Further mauve (Darling et al. 2004) was used for whole genome comparative studies. ClusatlX2 (Larkin et al. 2007), Mega7 (Kumar et al. 2016) and Dnasp v5 (Librado and Rozas 2009) were used for multiple sequence alignment, computation of pairwise distance and comparative sequence analysis. Phylogenetic tree was constructed by maximum likelihood method using 100 bootstrap replicates in MEGA7. Ka/Ks ration was calculated by DnaSPV5 and PamL4.9 (Xu and Yang 2013). SNPs and nucleotide diversity was analyzed by Mauve and DnaSPV5.

## Acknowledgements

Authors are thankful to UGC-CAS Department of Biosciences, Department of higher education, and State Government of Gujarat for the financial support. The first author is thankful to DST inspire fellowship program for providing research grants.

## Data Accessibility

The accession number of complete chloroplast genome of *Cressa cretica* deposited in Genbank accession: NC_035516 **(8th May 2017)**

## References

Bennett MD, Leitch IJ (1995) Nuclear DNA amounts in angiosperms. Ann Bot 76:113–176

Bock R (2007) Structure, function, and inheritance of plastid genomes. In: Cell and molecular biology of plastids. Springer, pp 29–63

Cai Z, Guisinger M, Kim H-G, Ruck E, Blazier JC, McMurtry V, Kuehl JV, Boore J, Jansen RK (2008) Extensive reorganization of the plastid genome of Trifolium subterraneum (Fabaceae) is associated with numerous repeated sequences and novel DNA insertions. J Mol Evol 67:696–704

Chen Z, Grover CE, Li P, Wang Y, Nie H, Zhao Y, Wang M, Liu F, Zhou Z, Wang X (2017) Molecular evolution of the plastid genome during diversification of the cotton genus. Mol Phylogenet Evol 112:268–276

Chumley TW, Palmer JD, Mower JP, Fourcade HM, Calie PJ, Boore JL, Jansen RK (2006) The complete chloroplast genome sequence of Pelargoniumx hortorum: organization and evolution of the largest and most highly rearranged chloroplast genome of land plants. Mol Biol Evol 23:2175–2190

Darling AC, Mau B, Blattner FR, Perna NT (2004) Mauve: multiple alignment of conserved genomic sequence with rearrangements. Genome Res 14:1394–1403

Downie SR, Jansen RK (2015) A comparative analysis of whole plastid genomes from the Apiales: expansion and contraction of the inverted repeat, mitochondrial to plastid transfer of DNA, and identification of highly divergent noncoding regions. Syst Bot 40:336–351

Gregory TR (2002) A bird’s-eye view of the C-value enigma: genome size, cell size, and metabolic rate in the class Aves. Evolution 56:121–130

Guisinger MM, Kuehl JV, Boore JL, Jansen RK (2011) Extreme reconfiguration of plastid genomes in the angiosperm family Geraniaceae: rearrangements, repeats, and codon usage. Mol Biol Evol 28:583–600

Guo W, Grewe F, Cobo-Clark A, Fan W, Duan Z, Adams RP, Schwarzbach AE, Mower JP (2014) Predominant and substoichiometric isomers of the plastid genome coexist within Juniperus plants and have shifted multiple times during cupressophyte evolution. Genome Biol Evol 6:580–590

Hirao T, Watanabe A, Kurita M, Kondo T, Takata K (2008) Complete nucleotide sequence of the Cryptomeria japonica D. Don. chloroplast genome and comparative chloroplast genomics: diversified genomic structure of coniferous species. BMC Plant Biol 8:70

Jansen RK, Raubeson LA, Boore JL, Chumley TW, Haberle RC, Wyman SK, Alverson AJ, Peery R, Herman SJ, Fourcade HM (2005) Methods for obtaining and analyzing whole chloroplast genome sequences. Methods in enzymology 395:348–384

Jockusch EL (1997) An evolutionary correlate of genome size change in plethodontid salamanders. Proc R Soc Lond B Biol Sci 264:597–604

Knight CA, Ackerly DD (2002) Variation in nuclear DNA content across environmental gradients: a quantile regression analysis. Ecol Lett 5:66–76

Kumar S, Stecher G, Tamura K (2016) MEGA7: Molecular Evolutionary Genetics Analysis version 7.0 for bigger datasets. Mol Biol Evol 18:1870–1874

Larkin MA, Blackshields G, Brown N, Chenna R, McGettigan PA, McWilliam H, Valentin F, Wallace IM, Wilm A, Lopez R (2007) Clustal W and Clustal X version 2.0. bioinformatics 23:2947–2948

Lee H-L, Jansen RK, Chumley TW, Kim K-J (2007) Gene relocations within chloroplast genomes of Jasminum and Menodora (Oleaceae) are due to multiple, overlapping inversions. Mol Biol Evol 24:1161–1180

Librado P, Rozas J (2009) DnaSP v5: a software for comprehensive analysis of DNA polymorphism data. Bioinformatics 25:1451–1452

Lin C-P, Huang J-P, Wu C-S, Hsu C-Y, Chaw S-M (2010) Comparative chloroplast genomics reveals the evolution of Pinaceae genera and subfamilies. Genome Biol Evol 2:504–517

Liu C, Shi L, Zhu Y, Chen H, Zhang J, Lin X, Guan X (2012) CpGAVAS, an integrated web server for the annotation, visualization, analysis, and GenBank submission of completely sequenced chloroplast genome sequences. BMC genomics 13:715

Lowe TM, Eddy SR (1997) tRNAscan-SE: a program for improved detection of transfer RNA genes in genomic sequence. Nucleic Acids Res 25:955–964

Luo J, Hou B-W, Niu Z-T, Liu W, Xue Q-Y, Ding X-Y (2014) Comparative chloroplast genomes of photosynthetic orchids: insights into evolution of the Orchidaceae and development of molecular markers for phylogenetic applications. PloS one 9:e99016

Magee AM, Aspinall S, Rice DW, Cusack BP, Sémon M, Perry AS, Stefanović S, Milbourne D, Barth S, Palmer JD (2010) Localized hypermutation and associated gene losses in legume chloroplast genomes. Genome Res 20:1700–1710

McNeal JR, Kuehl JV, Boore JL, de Pamphilis CW (2007) Complete plastid genome sequences suggest strong selection for retention of photosynthetic genes in the parasitic plant genus Cuscuta. BMC Plant Biol 7:57

Millen RS, Olmstead RG, Adams KL, Palmer JD, Lao NT, Heggie L, Kavanagh TA, Hibberd JM, Gray JC, Morden CW (2001) Many parallel losses of infA from chloroplast DNA during angiosperm evolution with multiple independent transfers to the nucleus. The Plant Cell 13:645–658

Milligan BG, Hampton JN, Palmer JD (1989) Dispersed repeats and structural reorganization in subclover chloroplast DNA. Mol Biol Evol 6:355–368

Morden CW, Wolfe K (1991) Plastid translation and transcription genes in a non-photosynthetic plant: intact, missing and pseudo genes. EMBO J 10:3281

Nevo E (2001) Evolution of genome-phenome diversity under environmental stress. Proc Natl Acad Sci 98:6233–6240

Palmer JD, Osorio B, Aldrich J, Thompson WF (1987) Chloroplast DNA evolution among legumes: loss of a large inverted repeat occurred prior to other sequence rearrangements. Curr Genet 11:275–286

Palmer JD, Thompson WF (1982) Chloroplast DNA rearrangements are more frequent when a large inverted repeat sequence is lost. Cell 29:537–550

Perry AS, Wolfe KH (2002) Nucleotide substitution rates in legume chloroplast DNA depend on the presence of the inverted repeat. J Mol Evol 55:501–508

Plunkett GM, Downie SR (2000) Expansion and contraction of the chloroplast inverted repeat in Apiaceae subfamily Apioideae. Syst Bot 25:648–667

Priyashree S, Jha S, Pattanayak S (2010) A review on Cressa cretica Linn.: A halophytic plant. Pharmacogn Rev 4:161

Raubeson LA, Jansen RK (1992) A rare chloroplast-DNA structural mutation is shared by all conifers. Biochem Syst Ecol 20:17–24

Raubeson LA, Jansen RK (2005) Plant diversity and evolution: genotypic and phenotypic variation in higher plants. Cabi Publishing

Saski C, Lee S-B, Daniell H, Wood TC, Tomkins J, Kim H-G, Jansen RK (2005) Complete chloroplast genome sequence of Glycine max and comparative analyses with other legume genomes. Plant Mol Biol 59:309–322

Schmitz-Linneweber C, Maier RM, Alcaraz J-P, Cottet A, Herrmann RG, Mache R (2001) The plastid chromosome of spinach (Spinacia oleracea): complete nucleotide sequence and gene organization. Plant Mol Biol 45:307–315

Shi C, Hu N, Huang H, Gao J, Zhao Y-J, Gao L-Z (2012) An improved chloroplast DNA extraction procedure for whole plastid genome sequencing. PLoS One 7:e31468

Tsudzuki J, Nakashima K, Tsudzuki T, Hiratsuka J, Shibata M, Wakasugi T, Sugiura M (1992) Chloroplast DNA of black pine retains a residual inverted repeat lacking rRNA genes: nucleotide sequences of trnQ, trnK, psbA, trnI and trnH and the absence of rps16. Molecular and General Genetics MGG 232:206–214

Turmel M, Otis C, Lemieux C (2005) The complete chloroplast DNA sequences of the charophycean green algae Staurastrum and Zygnema reveal that the chloroplast genome underwent extensive changes during the evolution of the Zygnematales. BMC Biol 3:22

Wang R-J, Cheng C-L, Chang C-C, Wu C-L, Su T-M, Chaw S-M (2008) Dynamics and evolution of the inverted repeat-large single copy junctions in the chloroplast genomes of monocots. BMC Evol Biol 8:36

Wicke S, Schäferhoff B, Müller KF (2014) Disproportional plastome-wide increase of substitution rates and relaxed purifying selection in genes of carnivorous Lentibulariaceae. Mol Biol Evol 31:529–545

Wicke S, Schneeweiss GM, Müller KF, Quandt D (2011) The evolution of the plastid chromosome in land plants: gene content, gene order, gene function. Plant Mol Biol 76:273–297

Wolf PG, Roper JM, Duffy AM (2010) The evolution of chloroplast genome structure in ferns. Genome 53:731–738

Wolfe KH, Mordent CW, Ems SC, Palmer JD (1992) Rapid evolution of the plastid translational apparatus in a nonphotosynthetic plant: loss or accelerated sequence evolution of tRNA and ribosomal protein genes. J Mol Evol 35:304–317

Wu C-S, Chaw S-M (2015) Evolutionary stasis in cycad plastomes and the first case of plastome GC-biased gene conversion. Genome Biol Evol 7:2000–2009

Wyman SK, Jansen RK, Boore JL (2004) Automatic annotation of organellar genomes with DOGMA. Bioinformatics 20:3252–3255

Xu B, Yang Z (2013) PAMLX: a graphical user interface for PAML. Mol Biol Evol 30:2723–2724

Yang Z (1998) Likelihood ratio tests for detecting positive selection and application to primate lysozyme evolution. Mol Biol Evol 15:568–573

Zhang H, Li C, Miao H, Xiong S (2013) Insights from the complete chloroplast genome into the evolution of Sesamum indicum L. PLoS One 8:e80508

Zhu A, Guo W, Gupta S, Fan W, Mower JP (2016) Evolutionary dynamics of the plastid inverted repeat: the effects of expansion, contraction, and loss on substitution rates. New Phytol 209:1747–1756

